## Supplementary figures and images for "*replicAnt*: A pipeline for generating annotated images of animals in complex environments using Unreal Engine"

### Appendix Figure 1

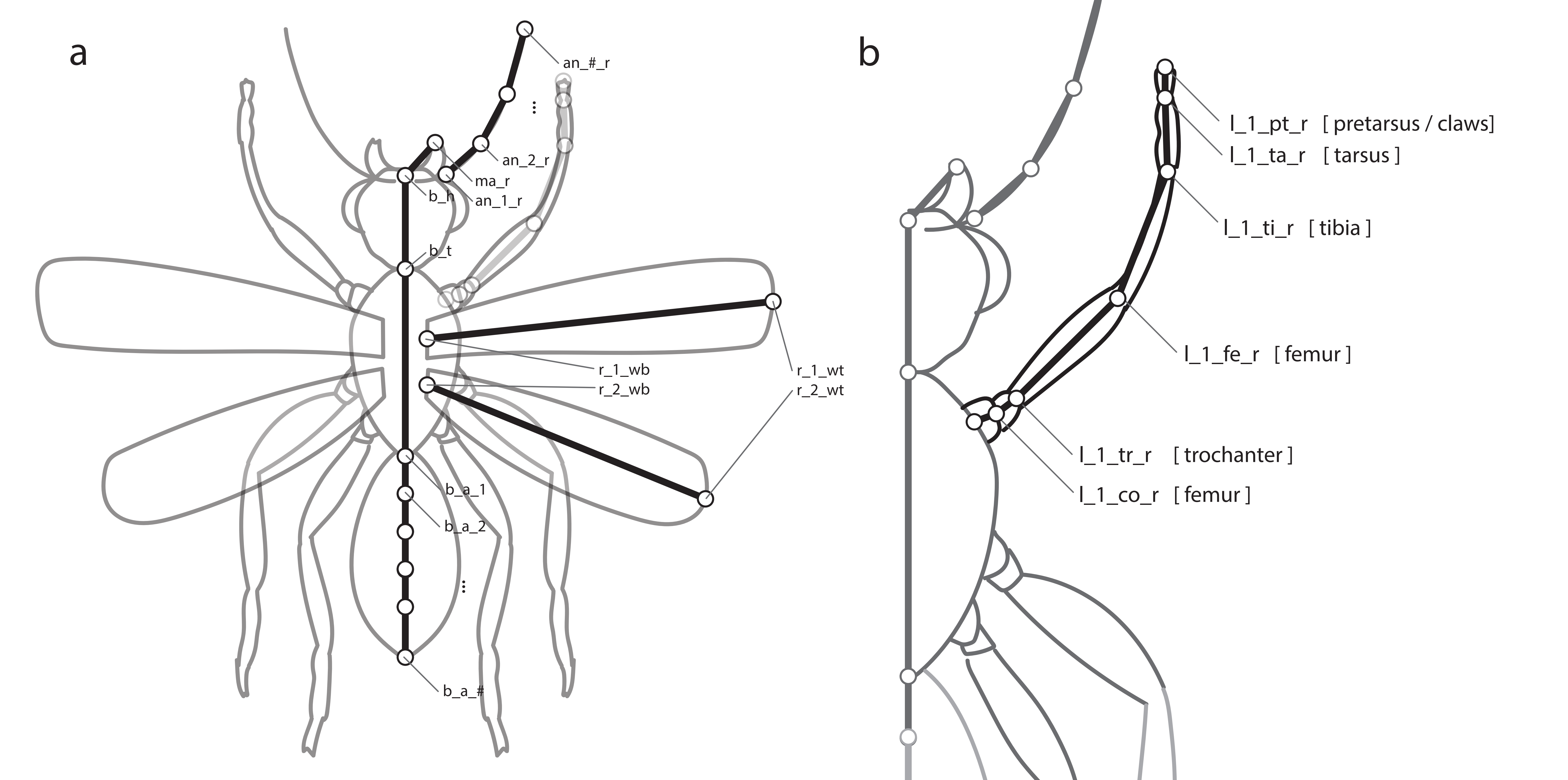
